# Localization of Realistic Spatial Patches of Complex Source Activity in MEG

**DOI:** 10.1101/2025.06.04.657819

**Authors:** Amita Giri, Lukas Hecker, John C. Mosher, Amir Adler, Dimitrios Pantazis

## Abstract

Accurate localization of neural sources in Magnetoencephalography (MEG) and Electroencephalography (EEG) is essential for advancing clinical and research applications in neuroscience. Traditional approaches like dipole fitting (e.g., MUSIC, RAP-MUSIC) are limited to discrete focal sources, while distributed source imaging methods (e.g., MNE, sLORETA) assume sources distributed across the cortical surface. These methods, however, often fail to capture sources with complex spatial extents, limiting their accuracy in realistic settings. To address these limitations, we introduce PATCH-AP, an enhanced version of the Alternating Projection (AP) method that effectively localizes both discrete and spatially extended sources. We evaluated PATCH-AP against leading source localization methods, including distributed source imaging techniques (MNE, sLORETA), traditional dipole fitting (AP), and recent extended source methods (Convexity-Champagne (CC), FLEX-AP). PATCH-AP consistently outperformed these methods in simulations, achieving lower Earth Mover’s Distance (EMD) scores—a metric indicating closer alignment with the true source distribution. In tests with real MEG data from a face perception task, PATCH-AP demonstrated high alignment with the fusiform face area, a region critical for face processing. These results highlight PATCH-AP’s potential to enhance source localization accuracy, promising significant advancements in neuroscience research and clinical diagnostics.

## I. Introduction

Magnetoencephalography (MEG) and electroencephalography (EEG) are widely utilized techniques known for their excellent temporal resolution, making them invaluable for capturing dynamic neural activity [1]–[6]. These techniques provide key insights into the intricate dynamics of both healthy and neurologically impaired brains. A significant drawback, however, of MEG and EEG is their limited spatial resolution, which presents challenges for accurately localizing neural sources. This difficulty stems from the inherently ill-posed nature of the inverse problem in source localization: the potential neural source locations far outnumber the sensors on or around the scalp, leading to an infinite number of possible source configurations that could explain the recorded signals.

Inverse source localization methods are broadly classified into two categories [7], [8]: distributed source imaging and dipole fitting. Distributed source imaging models assume sources are spread across the entire cortical surface, where source localization involves estimating a density map of these active sources using linear optimization techniques such as the minimum norm estimate (MNE) [9], weighted MNE (WMNE) [10], [11], low-resolution electromagnetic tomography (LORETA) [12], standardized LORETA (sLORETA) [13], and local autoregressive average (LAURA) [14]. Despite their widespread use, these methods often struggle to accurately identify sparse or focal sources.

Dipole fitting methods, in contrast, avoid the ill-posedness of the inverse problem by identifying a small set of equivalent current dipoles (ECDs) whose fields best match the M/EEG measurements using a least-squares approach [15]. Rather than assigning a current value to every possible dipole across a densely sampled source space, these methods concentrate on pinpointing specific focal sources. Key dipole fitting methods include beamformers [16], [17] and MUSIC [15], as well as recursive extensions such as RAP-MUSIC [18], Truncated RAP-MUSIC [19], and RAP Beamformer [20]. While recursive methods generally outperform non-recursive versions, they face limitations including a dependency on high signal-to-noise ratio (SNR) and the risk of canceling correlated sources. Recent advancements, such as the Alternating Projections (AP) method [21], [22] and DS-MUSIC [20], have addressed some of these challenges. In particular, the AP method has demonstrated reduced localization errors compared to RAP-MUSIC, especially in scenarios with high source correlation or low SNR, making it a highly effective option for real-world applications [21].

Although AP performs well in low SNR and high-correlation settings, it struggles to accurately model sources with spatial extent, which cannot be represented as single dipoles. To address this, the FLEX-AP method [23] was recently developed to extend AP by estimating both focal and extended sources through progressively smoothed matrices for source patches of increasing extent. FLEXAP, however, assumes full coherence (rank-1) for all components of an extended source, which does not always hold, as cortical patch activities often display varying degrees of coherence and may deviate from a rank-1 model. In summary, current methods tend to favor either discrete or distributed sources, and even when extended sources are addressed, complete coherence is generally assumed—an assumption that may not reflect realistic neural activity.

To overcome these limitations, we introduce PATCH-AP, an enhanced version of the AP method designed to localize both discrete and extended neural sources. PATCH-AP bridges the simplicity of dipole fitting with the comprehensiveness of distributed source imaging by effectively modeling and localizing both rank-1 focal sources with no spatial extent as well as rank-2 extended patches. PATCH-AP was evaluated against existing methods, including distributed source imaging (MNE, sLORETA), dipole fitting methods (AP), and recent advances in extended source localization (Convexity-Champagne (CC) and FLEX-AP). PATCH-AP demonstrated superior performance in simulations, as shown by lower Earth Mover’s Distance (EMD) values, which measure the minimum cost to align the estimated source distribution with the true source distribution. We assessed PATCH-AP’s performance under varying conditions, including signal-to-noise ratios (SNR), inter-source correlations (ISC), and patch smoothness orders, and validated its applicability using real human MEG data from a face perception task.

This study introduces the PATCH-AP method and evaluates its source localization performance relative to existing methods. Section II presents the problem formulation for localizing spatially smooth source signals, followed by the approach for introducing a rank-*r* approximation to model activation time courses. Section III details the computation of the PATCH-AP inverse solution. Section IV describe the methodology for estimating the amplitudes of the source patches. Results from simulated and real EEG data evaluations are provided in Section V. The article concludes with a discussion in Section VI. Our findings demonstrate that PATCHAP improves extended neural source localization accuracy, holding significant promise for advancing clinical and research applications in neuroscience.

## II. Materials & Methods

### A. Background

The electrical currents generated by the apical dendrites of pyramidal neurons in the cerebral cortex are generally considered the primary source of MEG and EEG signals [24], [25]. Clusters of thousands of pyramidal cortical neurons that are activated simultaneously can be represented as an equivalent current dipole (ECD), which serves as the fundamental unit for modeling neural activity in MEG and EEG localization techniques.

In M/EEG recordings, the data **y**(*t*) ∈ ℝ^*m*^ captured by *m* sensors at a given time *t* can be expressed as the sum of responses from *k* current dipoles distributed across the source space. This relationship is mathematically represented as:

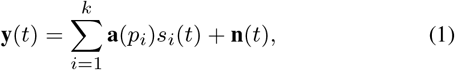

where **a**(*p*_*i*_) denotes the forward vector for the *i*th source at position *p*_*i*_, *s*_*i*_(*t*) represents the source activity at time *t*, and **n**(*t*) ∈ ℝ^*m*^ represents additive white Gaussian noise. The forward vector **a**(*p*_*i*_) is defined as:

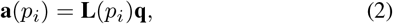

where **L**(*p*_*i*_) is the *m* × 3 forward matrix at location *p*_*i*_, and **q** is the 3 × 1 orientation vector of the ECD source. Depending on the application, the dipole orientation **q** may be fixed (known) or freely oriented (unknown and to be estimated) [3], [15].

Assuming that recordings are obtained at *T* discrete time points *t* ∈ [1, …, *T*], the matrix **Y** representing the sampled signals can be formulated as:

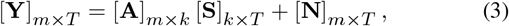

where **Y** is the *m* × *T* EEG/MEG data matrix:

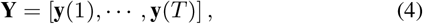

**A** is the *m* × *k* forward mapping matrix at the *k* candidate dipole locations comprising the entire source space:

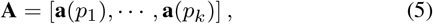

**S** is the *k* × *T* source time series matrix:

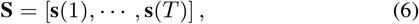

with **s**(*t*) = [*s*_1_(*t*), …, *s*_*k*_ (*t*)]^*T*^, and **N** is the *m* × *T* noise data matrix:

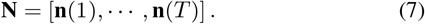

In this work, we restrict analyses to *fixed-orientated* dipoles, which align with the collective orientation of pyramidal cell dendrites [15], [24], [25]. Traditional dipole fitting methods, such as MUSIC, RAPMUSIC, and AP, estimate the positions of *N* focal active sources from a set of *k* dipoles representing the entire source space. These methods assume that source signals (**S**) arise from rank-1 dipole sources and do not account for extended sources. To address this limitation, the FLEX method [23] was developed to model extended brain sources, though it still assumes rank-1 sources.

Here, we propose the PATCH-AP method, which estimates extended sources, or “patches,” with a rank greater than 1, in contrast to prior methods that assume rank-1 sources. Notably, localizing both the positions and spatial extent of sources in patch models pose an inherently ill-posed inverse problem, as the number of unknown parameters (location and extent) exceeds the number of sensors available in EEG or MEG setups. To manage this complexity, we introduce source priors to constrain the source space: (1) spatial smoothness of sources, and (2) activation time courses with reduced rank.

### B. Spatially Smooth Source Signals

This section addresses the spatial smoothing of source signals, an essential property to account for correlations between neighboring dipoles within source patches. Let *I* ∈ [1, …, *N*] denote the index of the source patches. The set *P* = {*p*_1_, …, *p*_*N*_} represents the seed dipole (or center) locations of the source patches, and *L* = {*l*_1_, …, *l*_*N*_} denotes the corresponding source extents. The source extent *l*_*i*_ is an integer that defines the order of the patch extent, indicating the inclusion of neighboring dipoles up to a order of *l*_*i*_ from the center of the patch. For the *i*th patch, with location *p*_*i*_ and extent *l*_*i*_, the number of dipoles in the patch is given by *d*_*i*_. The total number of dipoles across all patches is then *d* = *d*_1_ + *d*_2_ +… + *d*_*N*_.

To apply spatial smoothing to the source signals 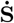 (of dimension *d* × *T*), we introduce a smoothing operator ***K***. This operator ***K***, which will rely on the Laplacian operator, is designed to smooth the signals across the cortical mesh, effectively incorporating the spatial structure of the sources within the model.

a. *Laplacian Operator on Cortical Mesh:* The cortical mesh is typically represented as a triangulated surface that spans the entire source space. Spatial relationships between sources are captured using an adjacency matrix 𝒜 ∈ ℝ^*k×k*^, where *k* denotes the number of grid points sampling the surface. This matrix encodes the spatial proximity of dipoles. Specifically, A is defined as:

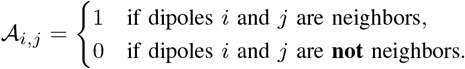

The adjacency matrix thus identifies which dipoles (sources) are spatially adjacent, allowing neighboring dipoles to influence each other’s signals during smoothing. The degree matrix ***D*** is defined as a diagonal matrix, where each diagonal element ***D***_*ii*_ is the sum of the elements in the *i*th row of the adjacency matrix:

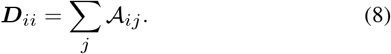

Using the above, the Laplacian matrix L is then computed as:

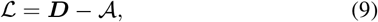

The Laplacian matrix captures local variations in the local neighborhood on the grid and plays a central role in modeling spatial smoothness on the cortex.
b. *Smoothing Operator:* The smoothing operator ***K*** is constructed using the Laplacian matrix and is defined as:

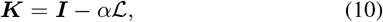

where 0 *< α <* 1 is a diffusion parameter that controls the degree of smoothing, and ***I*** is the identity matrix of size *k* × *k*.
c. *Patch Basis Matrix:* We define the patch basis matrix **E** ∈ ℝ ^*k×d*^ to represent the sources associated with each patch in the model. For each patch *i*, let 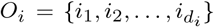 denote the set of indices corresponding to the sources within that patch. The vector 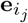 is a standard basis vector of dimension *k* × 1, with a value of 1 at the index *i*_*j*_ ∈ *O*_*i*_ and 0 elsewhere. Using these vectors, we construct the patch basis matrix:

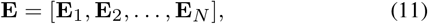

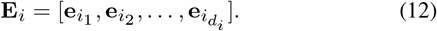

where *d*_*i*_ is the number of sources in patch *i*, and *N* is the total number of patches.
d. *Constructing Spatially Smooth Source Signals:* Finally, the spatially smooth source signals **S** are constructed by utilizing the above smoothing operator ***K*** with smoothness order *q* and the patch basis matrix **E**. Here, ***K***^*q*^ denotes the matrix ***K*** raised to the power *q*. The parameter *q* is an integer that determines the level of spatial smoothness, where *q* = 0 corresponds to no smoothing. To facilitate comprehension, we mark the dimensions of the individual elements:

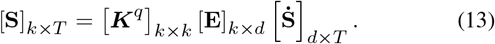

Using this, we can rewrite (3) as:

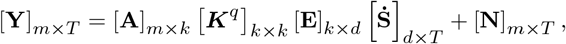

and to simplify notation in subsequent analysis:

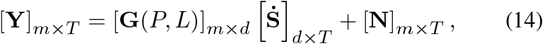

where

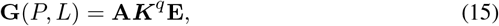

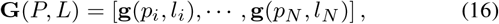

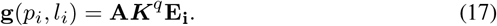

### C. Activation Time Courses of Reduced Rank

To make the source localization solution tractable, we approximate the forward model using a rank-*r* approximation. In the previous section, we described the forward model as:

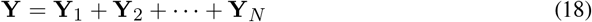

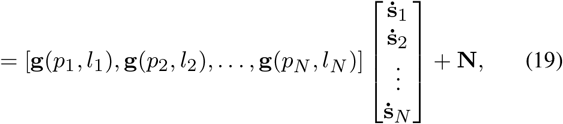

where **g**(*p*_*i*_, *l*_*i*_) represents the forward model for the *i*th source patch, and 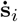 the corresponding source signal.

To reduce dimensionality, we perform a Singular Value Decomposition (SVD) of the forward model **g**(*p*_*i*_, *l*_*i*_) and approximate it with a rank-*r* approximation. The forward matrix for each source **g**(*p*_*i*_, *l*_*i*_) is decomposed as follows:

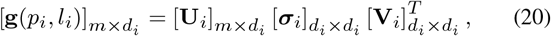

To approximate this matrix with reduced rank *r*_*i*_ (where *r*_*i*_ ≤ *d*_*i*_), we keep only the first *r*_*i*_ singular values and their corresponding singular vectors. This yields the rank-*r* approximation:

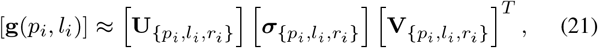

where:

- 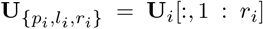 is the matrix of the first *r*_*i*_ left singular vectors,
- 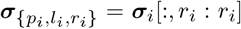 is the matrix of the first *r*_*i*_ singular values,
- 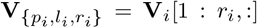 is the matrix of the first *r*_*i*_ right singular vectors.

Using the rank-*r* approximation of the forward matrix, we can rewrite the contribution of each source patch **Y**_*i*_ as:

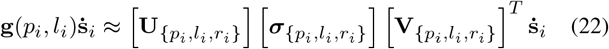

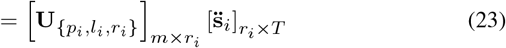

where 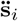 represents the reduced-rank approximation of the activation time courses for the *i*th patch.

During the source localization process, we estimate the reduced-rank *r*_*i*_ approximation of the activation time courses 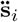, making the problem more tractable.

### D. Problem Formulation

In this section, we formulate the source localization problem, which aims to estimate the locations and extents of brain sources, as well as the corresponding source signals. The total signal recorded at *m* sensors can be written, utilizing the rank-reduced formulation from (23), as follows:

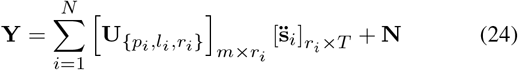

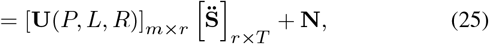

where *P* = {*p*_1_, …, *p*_*N*_} corresponds to the set of source locations, *L* = {*l*_1_, …, *l*_*N*_} is the set representing source extents, and *R* = {*r*_1_, …, *r*_*N*_} represents the set of ranks associated with each source. Here, *r* = *r*_1_ +*r*_2_ +… +*r*_*N*_ is the total rank of the system, combining contributions from all sources.

We define the objective function using a least-squares estimation criterion based on the observed sensor data **Y**. The problem can be stated as follows: *Given the observed data* **Y**, *estimate the N source locations P, their extents L, ranks R, and source time courses* 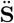. The least-squares estimation criterion is:

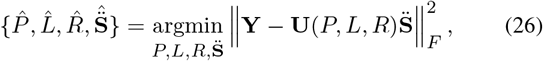

where **U**(*P, L, R*) represents the forward matrix parameterized by source positions *P*, extents *L*, and ranks *R*, and 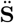 are the reduced-rank source activation time courses. The objective is to minimize the Frobenius norm of the difference between the observed sensor data **Y** and the modeled sensor data 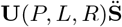.

This problem is further refined by introducing an assumption about the patch rank. In (10), we defined a smoothing operator that generates a sequence of forward matrices with progressively increasing smoothness. This operator is given by:

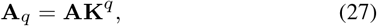

where **A** is the original forward matrix, and **A**_*q*_ is the smoothed forward matrix with smoothness order *q*. For each source patch, we compare the singular values of the original leadfield matrix to those of the smoothed leadfield matrix.

Figure 1 shows the cumulative percentage of energy captured by the singular value components. Notably, for as few as two singular components, the cumulative energy of the smoothed leadfield matrix exceeds 95%, effectively capturing most of the variance. This observation is supported by extensive Monte Carlo simulations of 500 patches with extents *l* ∈ [0, …, 5], where *l* = 0 represents a point source and *l* = 5 represents an extended source covering up to 5 neighboring dipoles (spanning 1.74 cm^2^ patch area on average). These simulations confirm that restricting the search to patches with a maximum rank of 2 is sufficient to capture the source contributions within each corresponding patch.

**Fig. 1.**
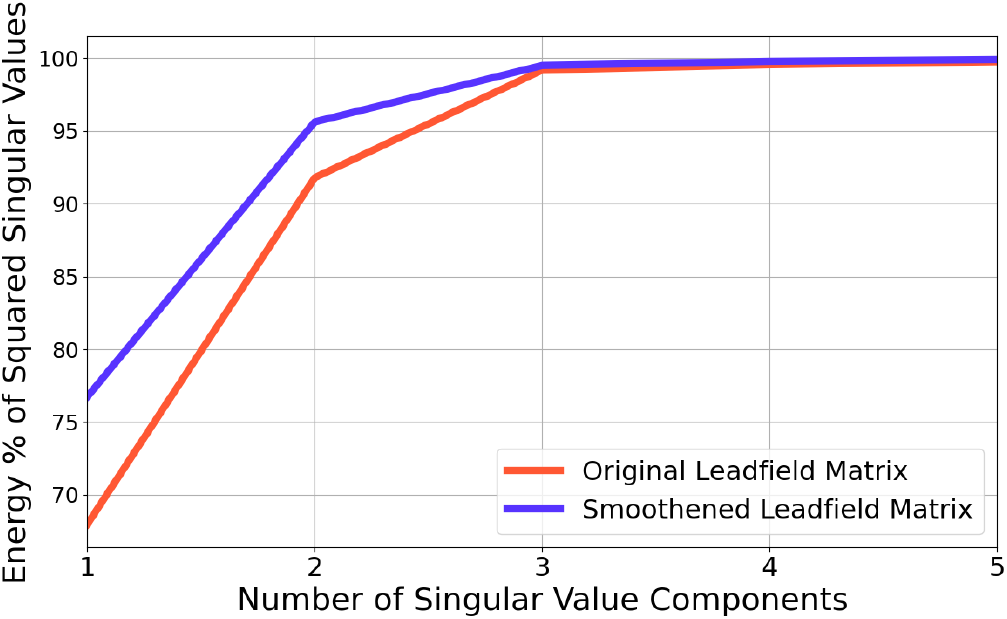
Energy Captured by Patch Activity of Different Ranks. Plot showing the cumulative energy percentage of squared singular values as a function of the number of singular value components for the original leadfield matrix (red) and the smoothed leadfield matrix (blue). The comparison highlights the effect of smoothing on the rank assumption.

Consequently, substituting *R* = {*r*_1_ = 2, …, *r*_*N*_ = 2} into (26), the objective function simplifies for the case of rank-2 sources:

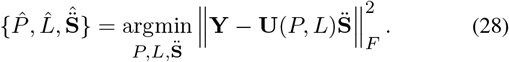

Thus, the estimated patch ranks are 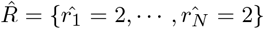. It is important to note that the rank is related to the estimated source extent. Specifically:

- For a point source (*l*_*i*_ = 0), the rank is *r*_*i*_ = 1.
- For an extended source (*l*_*i*_ *>* 0), the rank is set to *r*_*i*_ = 2.

This rank assumption allows the model to account for additional degrees of freedom in the activity of extended sources with non-zero extents. By incorporating this information, we enhance the localization accuracy of sources with varying spatial extents, effectively capturing their activation dynamics.

## III. PATCH-AP Inverse Solution

This section details the computation of the PATCH-AP inverse solution, which includes the standard Alternating Projections (AP) approach [21], [22] as a special case when the patch extent is set to 0 and the maximum rank is limited to 1. As noted previously, cortical patch activity is not fully coherent and does not always conform to rank 1. To address this limitation, we introduce an extension to the standard AP method, termed the PATCH-AP method.

In the earlier problem formulation section, the objective function for rank-2 sources was defined in (28). To simplify this further, we first express the unknown signal matrix 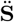 in terms of the modified forward matrix **U**(*P, L*). This leads to the following solution for 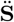:

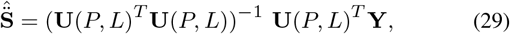

Defining the projection matrix **Π**_**U**(*P,L*)_ onto the column space of **U**(*P, L*) as:

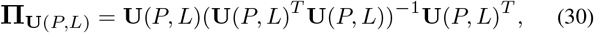

we can rewrite the objective function (28) as:

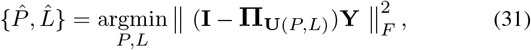

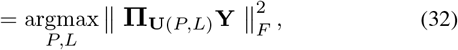

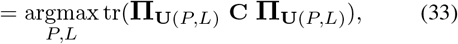

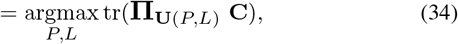

where *tr()* denotes the trace operator, **C** = **YY**^*T*^ is the data covariance matrix, and **I** is the identity matrix.

This formulation leads to a nonlinear and nonconvex *N* - dimensional maximization problem. The AP algorithm [26] addresses this by transforming the problem into a sequential, iterative process. During each iteration, the algorithm optimizes the parameters of a single source (location and extent) while holding the parameters of all other sources fixed at their previously estimated values.

Let *j* denote the current iteration and *n* the current source being estimated (with *n* = 1 to *N*). During this process, the parameters of other sources are fixed. For sources 1 to (*n* − 1), the parameters are 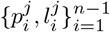, estimated in the current iteration. For sources *n* + 1 to *N*, the parameters 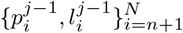 were estimated in the previous iteration.

The matrix 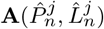 is defined as the *m* × (*N* − 1) matrix of topographies corresponding to these parameter values (excluding the *n*-th topography):

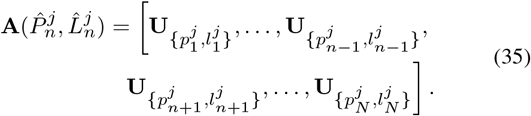

We decompose the projection matrix **Π** onto the column space of the augmented matrix 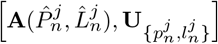 as:

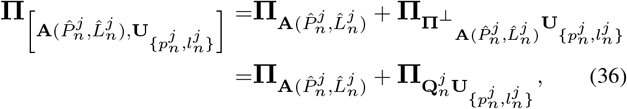

where

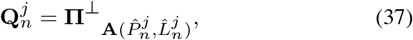

is the projection matrix that removes contributions from all sources except the *n*-th source in the *j*-th iteration.

Substituting (36) into (34) and neglecting the first term (as it does not depend on *p*_*n*_ and *l*_*n*_), we get:

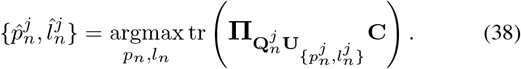

With:

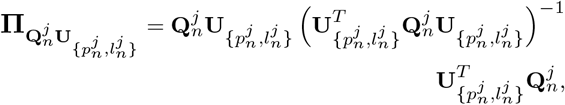

substituting this into equation (38), we finally obtain the PATCH-AP inverse solution:

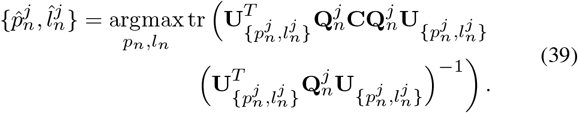

## IV. Patch Sources Amplitude Estimation

In this section, we describe the methodology for estimating the amplitudes of the source patches. For the *i*-th patch, the PATCH-AP method provides estimates of the patch’s location 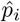, extent 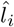, and rank 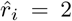. These parameters form the basis for estimating the source amplitudes **Ŝ** for each patch.

The first step is to compute 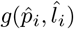 based on these estimated parameters, which describes the influence of each patch in the *m*-dimensional sensor space. We then apply SVD to 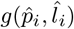, decomposing it into orthogonal components:

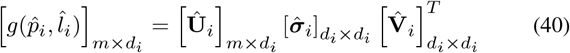

Here, **Û** _*i*_ and 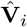 are orthogonal matrices representing the left and right singular vectors, respectively, and 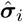 contains the singular values that describe the strength of the patch’s contribution to the sensor measurements. Since the estimated rank of each patch is 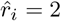, we approximate the contribution matrix 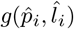 by retaining the top 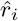 singular values. This rank reduction ensures computational feasibility while preserving the essential features of the patch’s contribution:

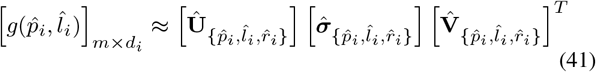

To estimate the source amplitudes 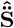 across all patches, we use a minimum-norm estimate (MNE) framework. For the *N* detected sources, with their estimated locations 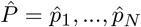, extents 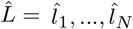, and ranks 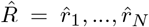, we form the combined left singular vector:

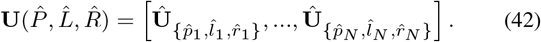

Since we assumed uniform rank, this simplifies to 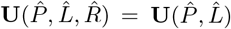.The amplitude of the sources within each patch can now be reconstructed by computing:

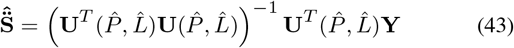

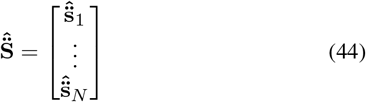

The source time course 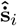 for each patch can be estimated using:

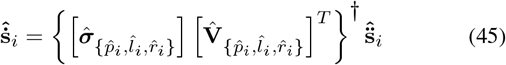

where † denotes the Moore-Penrose pseudoinverse. The overall source time courses for all patches can be combined into:

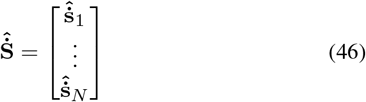

To achieve spatially smooth sources, the estimated source currents can be smoothed using a rank-specific spatial smoothing operator 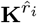 and estimated patch basis functions Ê as follows:

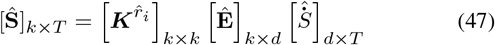

The process described ensures that the estimated source currents are not only rank-reduced but also spatially smooth and coherent with the underlying physical structure of the sources, making it more consistent with the realistic assumptions about the spatial distribution of the sources.

### A. Evaluation on Simulation

For our evaluation, we used an EEG setup with 128 channels arranged according to the International 10-20 system. To avoid the issue known as the “inverse crime”—which occurs when synthetic and inversion data share the same forward model assumptions—we employed distinct forward models for data synthesis and inversion [27]. Each simulation employed a source space comprising 5,124 fixed-oriented dipoles distributed across a cortical surface mesh. We set the conductivity ratio between the skull and brain to 1:80, using conductivities of 0.33 S/m^2^ for brain and scalp tissues and 0.0042 S/m^2^ for the skull [28]. The forward matrix was computed using the boundary element method (BEM) [29]. For inversion, we used the same forward matrix and two modified versions that avoid the inverse crime. The first variant increased the resolution to *k* = 8, 196 dipoles with octahedral spacing while retaining other parameters. The second variant maintained the higher resolution and adjusted the skull-to-brain conductivity ratio from 1:80 to 1:50.

We conducted an experiment to illustrate source localization results in multi-source scenarios with distinct rank combinations, alongside three additional experiments, each varying a single parameter and simulating 500 Monte Carlo samples of neural activity projected onto 128 EEG channels. Experiment 1 investigated the effects of varying noise levels by altering the Signal-to-Noise Ratio (SNR) to −5 dB, 0 dB, and 5 dB for a single patch of rank either [1] or [2], and a combination of two patches with ranks [1, 2]. The inter-source correlation was set to *ρ* = 0.5, and the smoothing parameter to 2. Experiment 2 examined the impact of inter-source correlation by adjusting *ρ* to {0.1, 0.5, 0.9} for patch ranks [1], [2] and [1,2]. We kept the SNR at 0 dB and used a smoothing parameter of 2. Experiment 3 analyzed the influence of smoothness order by varying it across 0, 2, 4 for patch ranks [1], [2] and [1,2], fixing the SNR at 0 dB, and the correlation coefficient at *ρ* = 0.5.

Inter-source correlation was introduced using a mixing matrix ***C***_*mix*_, derived from Cholesky decomposition:

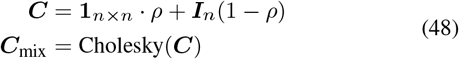

where ***I*** is the identity matrix, *ρ* represents the desired correlation, and *n* is the number of sources. The matrix ***C***_mix_ was then used to mix the source time courses. After projecting the source vectors ***J*** through the forward matrix, independent identically-distributed noise was added to achieve an SNR of 0 dB.

We evaluated the performance of the Patch-AP method against its greedy counterparts (AP and FLEX-AP) as well as established methods such as MNE, sLORETA, and Convexity-Champagne. FLEXAP was implemented following the approach outlined in Hecker et al. [23], [30]. For non-greedy methods, Tikhonov regularization was applied, with the optimal parameter selected from a set of ten options based on the generalized cross-validation criterion. For Convexity-Champagne, hyperparameters were tuned until the loss converged to a value below 10^−8^, using an active set strategy to eliminate near-zero candidates.

For the evaluation, we chose to use the Earth Mover’s Distance (EMD), also known as the Wasserstein metric. EMD quantifies the minimum effort required to transform one distribution into another, measured in millimeters based on the Euclidean distance between dipoles. The EMD calculations were performed using the Python POT package [31].

Figure 2 illustrates the rank-2 extended simulated source activity and the corresponding inverse solutions. Panel (a) shows the simulation of a rank-2 source at 0 dB SNR with no modeling error. Panels (b), (c), and (d) display the inverse solutions obtained using the MNE, sLORETA, and CC methods, respectively. Panel (e) presents the EMD metrics for these three inverse methods. Panels (f), (g), and (h) depict the inverse solutions from the AP, FLEX-AP, and PATCH-AP methods, respectively, while panel (i) shows the EMD metrics corresponding to the inverse solutions from AP, FLEX-AP, and PATCH-AP. This figure highlights the limitations of dipole imaging and dipole fitting methods when applied to sources that are neither strictly focal nor extended across the entire cortical surface. Notably, PATCH-AP demonstrates the lowest EMD among all evaluated methods, indicating superior accuracy in source localization.

**Fig. 2.**
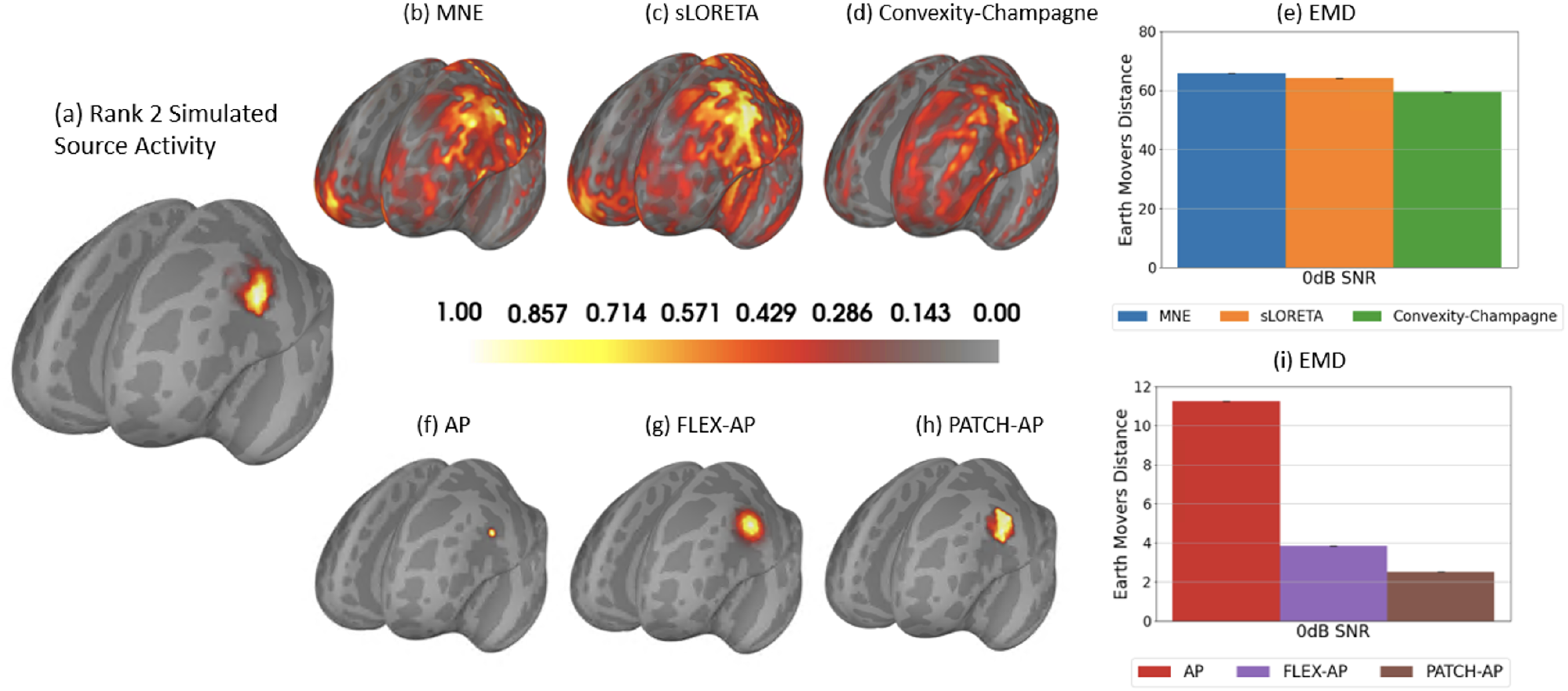
Simulated Extended Source Activity of Rank 2 and Corresponding Inverse Solutions. (a) Simulation of one rank 2 source at 0 dB SNR with no modeling error. (b) MNE inverse solution. (C) sLORETA inverse solution. (d) Convexity-Champagne inverse solution. (e) Earth mover’s Distance (EMD) corresponding to (b), (c) and (e) inverse methods. (f) AP inverse solution. (g) FLEX-AP inverse solution. (h) Patch-AP inverse solution. (i) Earth mover’s Distance (EMD) corresponding to (f), (g) and (h) inverse methods.

### B. Evaluation on Real Data

We evaluated the performance of the Patch-AP algorithm on the SPM Face Dataset, collected by R. Henson and available through the Statistical Parametric Mapping (SPM) toolbox ^1^. This dataset consists of two sessions in which a subject views 86 face images and 86 scrambled face images. The MEG data were recorded on a 275-channel CTF/VSM system with second-order axial gradiometers and synthetic third gradient denoising, sampled at 480 Hz. Due to one faulty sensor, 274 MEG channels were used in this dataset. For our analysis, we used the second session of MEG data to compare the performance of the Patch-AP method against traditional methods, including MNE, sLORETA, CC, AP, and FLEX-AP.

The raw MEG data were band-pass filtered with a high-frequency cutoff of 30 Hz using a finite impulse response (FIR) filter, designed with the “firwin” method, known for its flexibility and effectiveness in creating a windowed sinc FIR filter. This filtering step removes high-frequency noise and artifacts, retaining only the relevant frequency components for analysis. Next, the continuous MEG data were segmented into epochs to isolate neural responses related to specific events. A total of 168 epochs were extracted, each spanning from −0.2 seconds before to 0.6 seconds after the event onset. Baseline correction was applied using the interval from −0.2 to 0 seconds, normalizing the data by removing any DC offset present in the pre-stimulus period. These epochs were then averaged to produce evoked responses, and a contrast was computed between the two conditions: “Faces” versus “Scrambled Faces”.

After epoching, noise covariance was calculated using all 168 trials with the *mne*.*compute covariance* function, focusing on the pre-stimulus period (tmax=0) to capture only noise-related activity. An empirical method was used for the covariance estimation, and no rank constraint was imposed (rank=None), allowing the full noise spectrum to be included in subsequent analyses. The covariance matrix was then applied to whiten the evoked response and transform the forward matrix accordingly. The source model comprised *k* = 8196 vertices with fixed orientations along a cortical mesh.

## V. Results

We present four distinct experiments using simulated data to compare the accuracy of Patch-AP with other algorithms, evaluating performance under varying conditions: (A) Illustration in multi-source scenarios with distinct rank combination [1,3], (B) signal-to-noise ratios (SNRs), (C) inter-source correlation (ISC), and (D) smoothness order. Additionally, we analyze PATCH-AP inverse solutions applied to real SPM Faces data.

### A. Illustration in Multi-Source Scenarios with Distinct Rank Combinations

In this evaluation, we illustrate the inverse solution of PATCH-AP in comparison to other methods for scenarios involving multiple sources with distinct rank combinations. Figure 3 (a) illustrates one rank 1 and one rank 3 extended simulated source activity at 0 dB SNR with no modeling error. Both sources have an extent of 2, spanning a 0.26 cm^2^ patch area on the cortex. Figure 3 (b), (c), and (d) display the inverse solutions obtained using AP, FLEX-AP, and PATCH-AP methods, respectively. finally, 2 (e) shows the EMD metrics corresponding to the inverse solutions. The EMD results demonstrate the superior performance of the PATCH-AP method in accurately localizing sources in multi-source scenarios with distinct rank combinations, outperforming both the FLEX and AP methods. These findings emphasize the effectiveness of PATCH-AP in extended source localization and its potential for improved accuracy in complex source configurations.

**Fig. 3.**
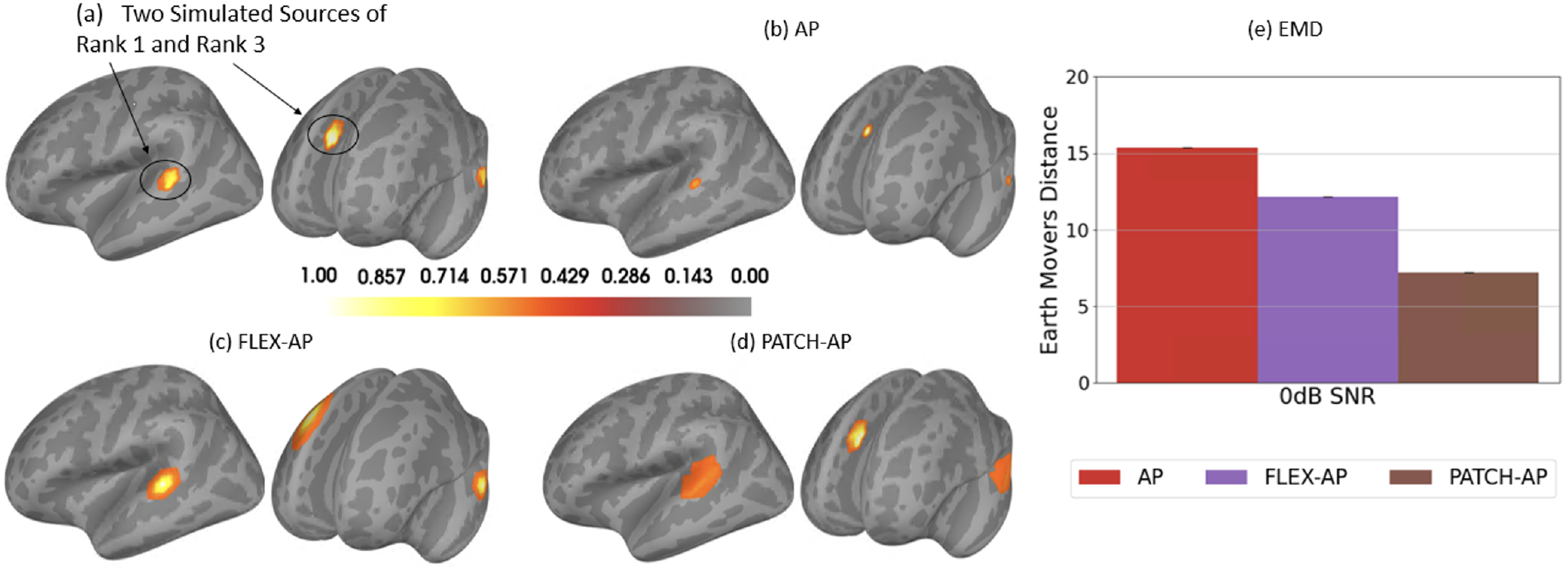
Multi-Source Scenarios with Distinct Rank Combinations. (a) Simulation of two sources with different ranks: one rank-1 source and one rank-3 source, at 0 dB SNR, with no modeling error. (b) AP inverse solution. (c) FLEX-AP inverse solution. (d) Patch-AP inverse solution. (e) Earth mover’s Distance (EMD) corresponding to (a), (b) and (c) inverse methods.

### B. Effect of SNR

The simulation experiment assessed the performance of the inverse solution under varying noise levels, simulating realistic MEG recordings with SNRs ranging from −5 to 5 dB. The Earth Mover’s Distance (EMD) metric was computed for different source configurations: a single source with a rank of [2], two sources with a rank combination of [1,2], and three sources with a rank combination of [1,2,2]. Figure 4 shows that PATCH-AP consistently outperformed MNE, sLORETA, CC, AP, and FLEX-AP across all SNRs and ranks. Notably, AP, FLEX-AP, and PATCH-AP demonstrated significantly better performance than MNE, sLORETA, and CC. PATCH-AP showed a clear advantage over MNE, sLORETA, and CC, while maintaining competitive performance with AP and FLEX-AP. Overall, the improvement in localization accuracy, as measured by EMD, was most pronounced with PATCH-AP.

**Fig. 4.**
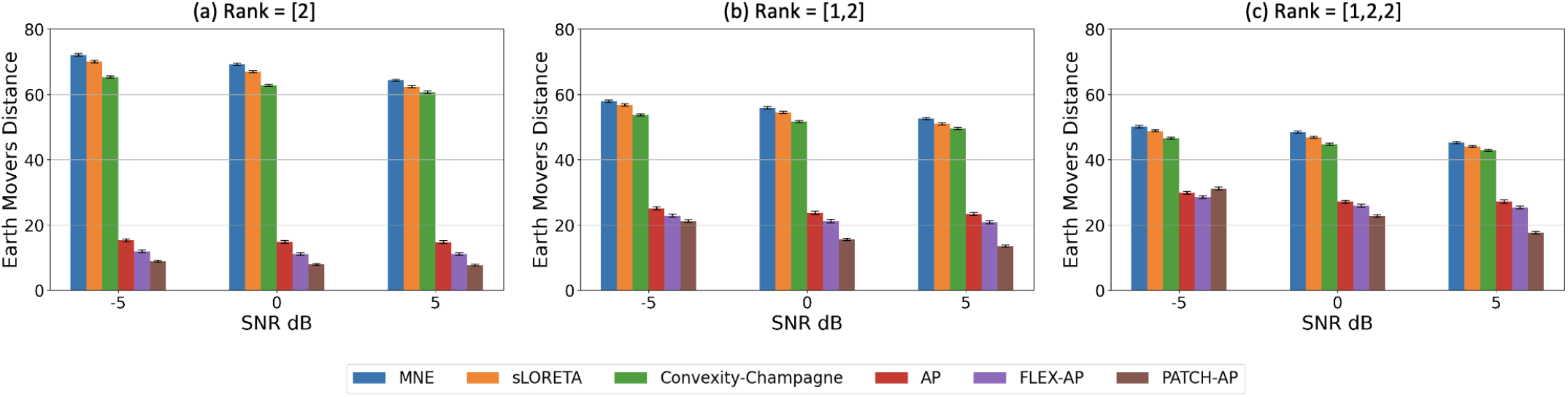
Effect of SNR on Inverse Solution Accuracy. Bar plots of the Earth Mover’s Distance (EMD) for various inverse solution methods at three SNR levels (−5 dB, 0 dB, and 5 dB), demonstrating the impact of noise on source localization accuracy. Results are presented for different source rank configurations: (a) Rank = [2], (b) Rank = [1, 2], and (c) Rank = [1, 2, 2]. The inter-source correlation coefficient is fixed at ***ρ* = 0.5**, and the smoothing parameter set to **2**. Lower EMD values indicate better localization performance, with the PATCH-AP method consistently showing superior results across all scenarios.

### C. Effect of ISC

Accurately identifying underlying sources becomes increasingly challenging when they are highly correlated, highlighting the need for methods that can effectively differentiate sources in such scenarios. Figure 6 illustrates the accuracy of PATCH-AP across a range of inter-source correlations, from 0.1 (low correlation) to 1 (complete coherence). The simulations included different source configurations: a single source with a rank of [2], two sources with a rank combination of [1,2], and three sources with a rank combination of [1,2,2]. AP, FLEX-AP, and PATCH-AP consistently outperformed MNE, sLORETA, and CC under both low and high correlation conditions. Among all methods, PATCH-AP achieved the lowest EMD across various correlation levels, demonstrating its robustness in accurately localizing sources even in the presence of source correlations.

**Fig. 5.**
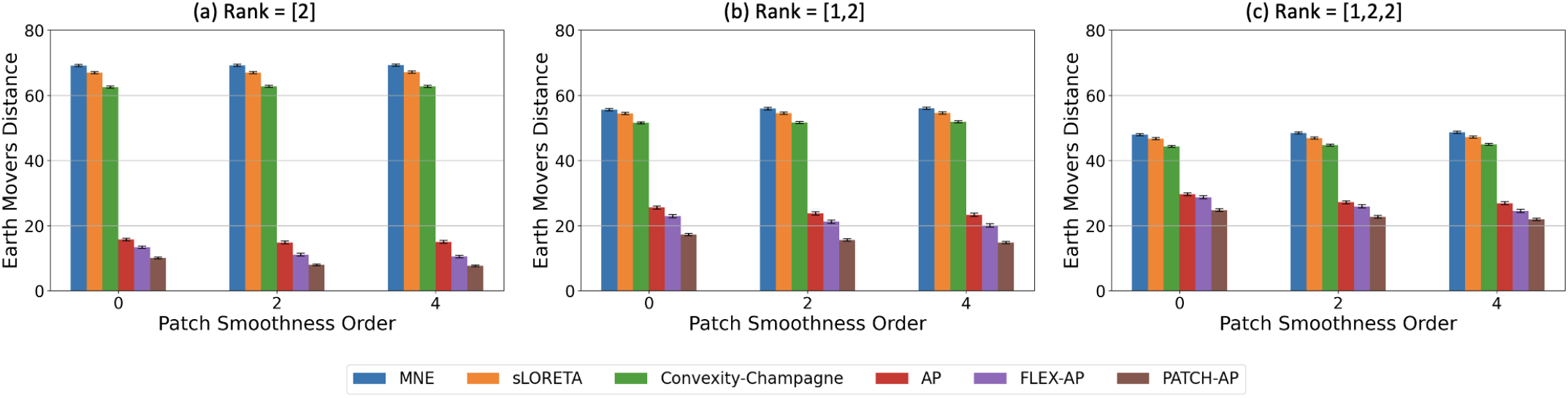
Effect of Patch Smoothness Order on Inverse Solution Accuracy. Earth Mover’s Distance (EMD) for various inverse methods across different patch smoothness orders (0, 2, 4). Results are shown for rank configurations: (a) Rank = [2], (b) Rank = [1, 2], and (c) Rank = [1, 2, 2], with simulations performed at 0 dB SNR and a correlation coefficient of ***ρ* = 0.5**.

**Fig. 6.**
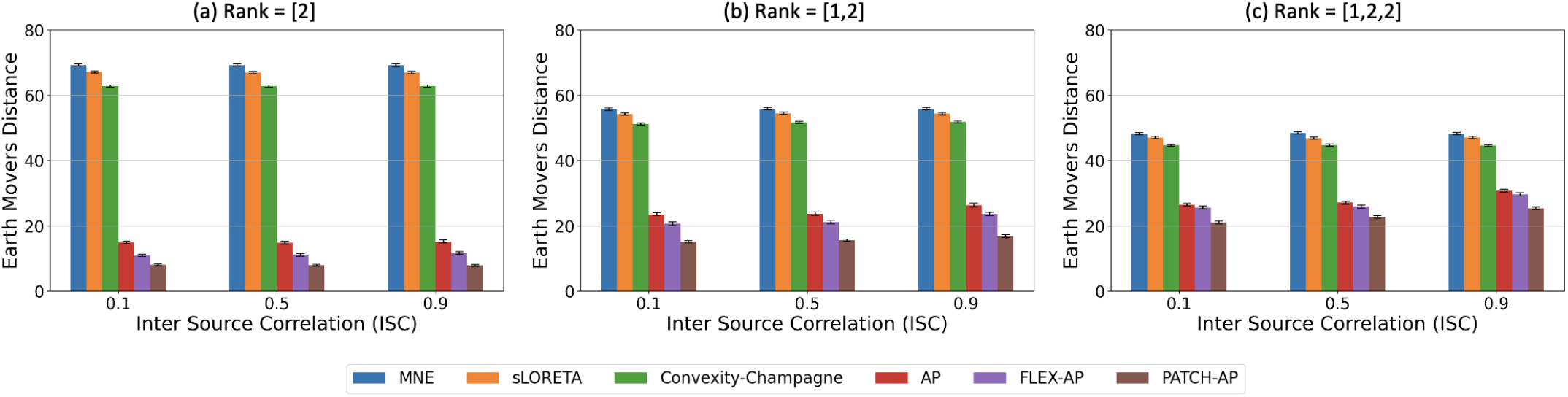
Effect of Inter-Source Correlation (ISC) on Inverse Solution Accuracy. Earth Mover’s Distance (EMD) for various inverse methods at different ISC levels (0.1, 0.5, 0.9), with results shown for rank configurations: (a) Rank = [2], (b) Rank = [1, 2], and (c) Rank = [1, 2, 2]. Simulations are performed at 0 dB SNR with a smoothing parameter of 2.

### D. Effect of Smoothness Order

In our simulations, we analyzed the performance of source localization methods across different patch smoothness orders. Figure 5 illustrates the EMD metric as the smoothness order varies from 0 (no smoothness) to 4 (high smoothness). The simulations included several source configurations: a single source with a rank of [2], two sources with a rank combination of [1,2], and three sources with a rank combination of [1,2,2]. The results highlight the superior performance of PATCH-AP compared to all other methods across various smoothness orders.

### E. SPM Real face data result

Validating these algorithms on real data, however, is essential to confirm their practical applicability. In this study, we applied the source localization methods to human MEG data recorded during a face perception task. Figure 7(a) illustrates the event-related fields (ERFs) in response to a contrast between faces and scrambled faces, averaged over 168 clean epochs. Our analysis focused on the time window from 140 ms to 190 ms (25 time samples) to compute inverse solutions using PATCH-AP and other established algorithms.

**Fig. 7.**
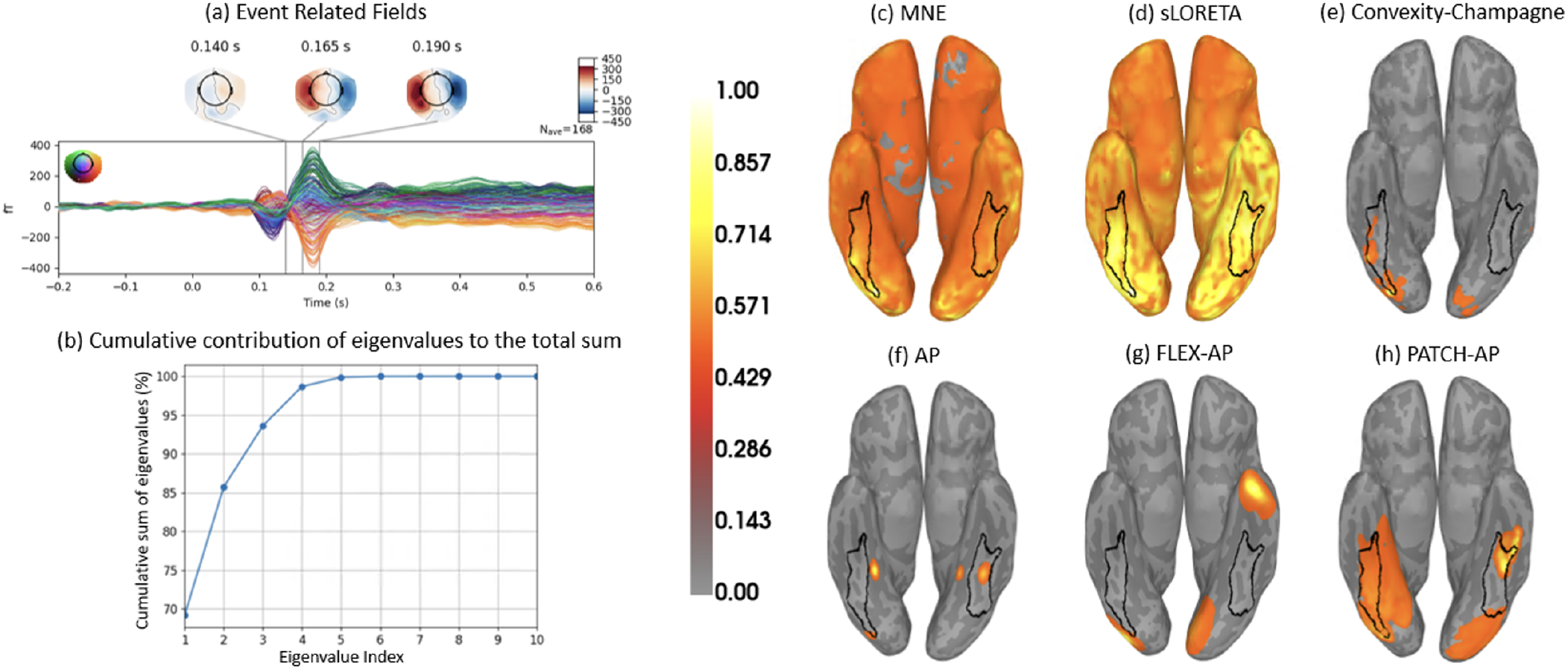
Source Localization of the SPM Faces Dataset. (a) Event-related fields (ERFs) in response to faces and scrambled objects, averaged over 168 trials and displayed as topographic maps of the magnetic fields in sensor space. (b) Cumulative contribution of eigenvalues to the total sum, showing the cumulative percentage of the total eigenvalue sum as a function of the eigenvalue index, indicating the number of sources required to capture significant variance. (c) MNE, (d) sLORETA, (e) Convexity-Champagne, (f) AP, (g) FLEX-AP, and (h) PATCH-AP inverse solutions. The PATCH-AP method demonstrates superior alignment in the fusiform face area (FFA), a region implicated in face processing, whereas AP and FLEX-AP exhibit lower localization accuracy.

To determine the number of sources in the real data, we assessed the ratio of the cumulative sum of eigenvalues to the total sum of the eigenvalues in the signal covariance matrix **C**. Figure 7(b) indicates the presence of four sources, as the cumulative sum surpasses 95% and eventually reaches 100%. Adding more sources beyond this point does not substantially increase the cumulative sum.

We then computed and visualized the source localization results, shown in Figures 7(c) through 7(h): (c) MNE, (d) sLORETA, (e) CC, (f) AP, (g) FLEX-AP, and (h) PATCH-AP. The proposed PATCH-AP method exhibits superior alignment with the fusiform face area (FFA), which is responsible for face processing, as highlighted by the black outline in the plot. In contrast, methods such as MNE, CC, AP, and FLEX-AP show less overlap with the FFA. Although sLORETA performs relatively well, showing an overlap comparable to PATCH-AP, the PATCH-AP method appears to offer the most accurate localization for both real and simulated data.

## VI. Conclusion

In summary, this study introduced PATCH-AP, an enhanced version of the Alternating Projection (AP) method, designed to overcome the limitations of traditional dipole fitting and distributed source imaging techniques in MEG source localization. PATCH-AP effectively localizes both discrete and extended sources, providing a more accurate representation of complex neural activity. Simulations demonstrated PATCH-AP’s robustness under varying SNR ratios, inter-source correlations, and patch smoothness conditions, consistently achieving lower EMD metrics compared to established methods like MNE, sLORETA, Convexity Champagne, AP, and FLEX-AP. Additionally, validation on real MEG data from a face perception task highlighted PATCH-AP’s superior alignment with the fusiform face area (FFA), confirming its practical utility in real-world applications. These results underscore PATCH-AP’s potential to significantly enhance clinical and research applications in neuroscience by enabling more precise and reliable neural source localization. Future research should continue to explore its efficacy across a broader range of experimental conditions and clinical scenarios, further establishing its value in neuroscience and clinical research.

## VII. Acknowledgements

This work was supported by the United States-Israel Binational Science Foundation grant 2020805 to A.A., and was supported in part by the National Eye Institute of the National Institutes of Health under award R01EY033638 to D.P., and by the National Institute on Aging of the National Institutes of Health under awards RF1AG074204 and RF1AG079324 to J.M. and D.P.. The content is solely the responsibility of the authors and does not necessarily represent the official views of the National Institutes of Health. The authors have no relevant personal financial or non-financial interests to disclose.

## Footnotes

1 https://www.fil.ion.ucl.ac.uk/spm/data/mmfaces/

